# Shifts in the gut microbiota structure caused by *Helicobacter pylori* eradication therapy

**DOI:** 10.1101/296426

**Authors:** Evgenii I. Olekhnovich, Alexander I. Manolov, Nikita A. Prianichniikov, Andrei E. Samoilov, Maja V. Malakhova, Alexander V. Pavlenko, Vlad V. Babenko, Andrei K. Larin, Yuriy Y. Babin, Elizaveta V. Starikova, Dmitry I. Chuvelev, Boris A. Kovarsky, Maria A. Tregubova, Dilyara D. Safina, Maria I. Markelova, Tatiana V. Grigoryeva, Eugenia A. Boulygina, Sergey Yu. Malanin, Rustam A. Abdulkhakov, Sayar R. Abdulkhakov, Elena S. Kostryukova, Elena N. Ilina, Vadim M. Govorun

**Affiliations:** Federal Research and Clinical Centre of Physical-Chemical Medicine, Malaya Pirogovskaya 1a, Moscow 119435, Russia; Kazan Federal University, 18 Kremlyovskaya, Kazan, 420008, Russia; Kazan State Medical University, 49 Butlerova, Kazan, 420012, Russia

## Abstract

The human gut microbiome plays an important role both in health and disease. The use of antibiotics can alter gut microbiota composition, which can cause complications of various kinds. Here we report a whole genome sequencing metagenomic study of the intestinal microbiota changes caused by *Helicobacter pylori* eradication therapy. We have found the decrease in taxonomic alpha-diversity due to the therapy. The changes observed were more extensive for patients with duodenal ulcer and female ones. As well across the patients under the therapy we have detected the shifts in the metabolic potential and resistome. Seven KEGG pathways associated with quorum sensing, genetic Information processing and environmental Information processing were increased, while metabolic pathways related with metabolism of cofactors and vitamins and glycan biosynthesis and metabolism decreased. Changes in the resistome profile have also been identified. We observed perturbations in intraspecies structures, which were higher in group of patients under the therapy than in control group of people without treatment. The *Eubacterium rectale* pangenome extracted from metagenomic data were changed. We also isolated and sequenced *Enterococcus faecium* strains from two patients before and after eradication therapy. After the therapy this bacterium increased as the antibiotic resistance in vitro, as well the number of ARGs to macrolides and tetracyclines and metagenomic relative abundance in comparison with strains before therapy. In summary, microbial community demonstrated shift to reduce metabolic potential and to increased mechanisms, which mediate more survival condition through intraspecies perturbations.

**Importance:** The human gut microbiome plays an important role both in health and disease. The use of antibiotics can alter gut microbiota composition, which can cause complications of various kinds. *H. pylori* eradication therapy causes multiple shifts and alterations (including intraspecies changes) of the intestinal microbiota structure and leads to the accumulation of genes which determine resistance to macrolides. Since these changes are not the same for patients with various diseases, patients with duodenal ulcer may be further paid special attention for reducing side effects, such as antibiotic-induced dysbiosis. Also, study of antibiotic treatment in terms of its impact upon the human gut microbiota allows shedding light on of the complex processes that cause accumulation and spread of antibiotic resistance. An identification and understanding of these complicated processes may help to constrain antibiotic resistance spread, which is of great importance for human health care.

## Introduction

*Helicobacter pylori* are ubiquitous bacteria classified by WHO as a cancer risk factor. It has been estimated that up to 50% of human population harbor this pathogen. It is supposed that *H. pylori* colonized the human stomach about 60 000 years ago after humanity’s ancestors migrated from Eastern Africa. Currently, *H. pylori* infection is associated with wide range of gastrointestinal diseases, including chronic gastritis, stomach ulcer, duodenal ulcer, gastric cancer, lymphoma of mucin-associated lymphoid tissue (Sanders et al., 2002). This bacterium is also responsible for iron-deficiency anemia, Werlhof disease in children (Queiroz et al., 2013), nutritional anemia due to vitamin B_12_ deficiency, thrombocytopenic purpura, lipid, and glucose metabolism disorders (Buzás, 2014).

Conventional *H. pylori* eradication therapy regimens are based on antibiotics (clarithromycin, amoxicillin, tetracycline, metronidazole), proton pump inhibitors and some other auxiliary drugs treatment, including bismuth salts (Malfertheiner et al., 2017). It is known that antibiotic treatment can significantly alter gut microbial community composition, and, as the result, may cause such complications as antibiotic-associated diarrhea (Bartlett et al., 2002; Tong et al., 2007). Bacteria can acquire antibiotic resistance by a number of ways that are associated either with single-nucleotide polymorphisms (SNPs) in antibiotic-target sites or with the exchange of the genes encoding efflux pumps, drug modifiers or proteins which protect antibiotic targets (Wright 2007; Davies and Davies 2010). Antibiotic resistance genes (ARGs) may be transferred among the bacteria by mobile genetic elements, bacteriophages and also as the result of natural transformation (Davies and Davies 2010; Schjørring and Krogfeldt 2011; Smillie et al. 2011). The term “resistome” has been coined to refer to the collection of ARGs in a certain microbial objects (Wright 2007; Marshall and Levy 2011). Resistome of human gut microbiota differs widely among the people of different countries and reflects peculiarities of antibiotic use (Forslund et al., 2013; Yarygin et al., 2017). Eradication therapy regimes include proton-pump inhibitors treatment which may lead to a significant shift in the gut microbial community due to the intrusion by the species that under normal conditions can’t pass gastric acid barrier (Jackson et al., 2016; Imhann et al., 2016).

Thus, eradication therapy can have a great impact on the gut microbial community composition. These changes can drive gut disorders and cause accumulation of ARGs in the gut microbial community. Here we used whole genome shotgun sequencing (WGS) to study taxonomic and functional profile shifts of the gut microbial community in the patients after *H. pylori* eradication therapy. Our research may contribute to better understanding of the processes in the gut microbiota, which are driven by the complex mix of drugs used during *H. pylori* eradication therapy.

## Results

### Cohorts of patients and available data

In total, 80 gut metagenomic samples collected from 40 patients (47.7±13.11 years old, 18 female and 22 male) before and after the *H. pylori* eradication therapy lasting 14 days were examined. The samples obtained were divided into groups to perform statistical analysis. The first group included metagenomes collected prior to the eradication therapy (1st time point, 40 metagenomic samples). The second group included metagenomes from samples collected 0-2 days after the end of therapy (2nd time point, 40 metagenomic samples).

The sequencing of the stool samples yielded 30.2±10.0 M of 50 bp reads per sample (120.9 Gbp in total). According to the rarefaction analysis results (Supplementary Figure S1), the obtained number of reads is sufficient to determine a significant part of the taxonomic and functional profile of the patient’s microbiota. Summary data on samples and patients are presented in additional data (Supplementary Tables S1).

### *H. pylori* eradication therapy affects composition and richness of gut microbiota

As a result of taxonomic profiling with MetaPhlAn2, 51 genus and 106 species were detected in the patients’ metagenomes. Eubacterium (15.46%±%13.45), Bacteroides (14.52%±16.92%) and Prevotella (14.43%±22.37%) were the most abundant genera at the 1st time point. The taxonomic composition of the 2nd point metagenomes was different: the most abundant genera included Bacteroides (22.77%±22.06%), Prevotella (14.06%±23.34%), Eubacterium (10.01%±16.08%). An increased relative abundance of Escherichia and Enterococcus was found after the therapy in the samples collected from four patients. Tables of taxonomic abundances at the genera and species levels are presented in additional materials (Supplementary Tables S2-S3). The GraPhlAn visualization of the patient’s gut microbiota community for two time points is presented in the Supplementary Figure S2. The differences in the taxonomic structure of the patient’s intestinal microbiota at two time points according to LEfSe analysis are presented in Figure 1.

**Figure 1.**
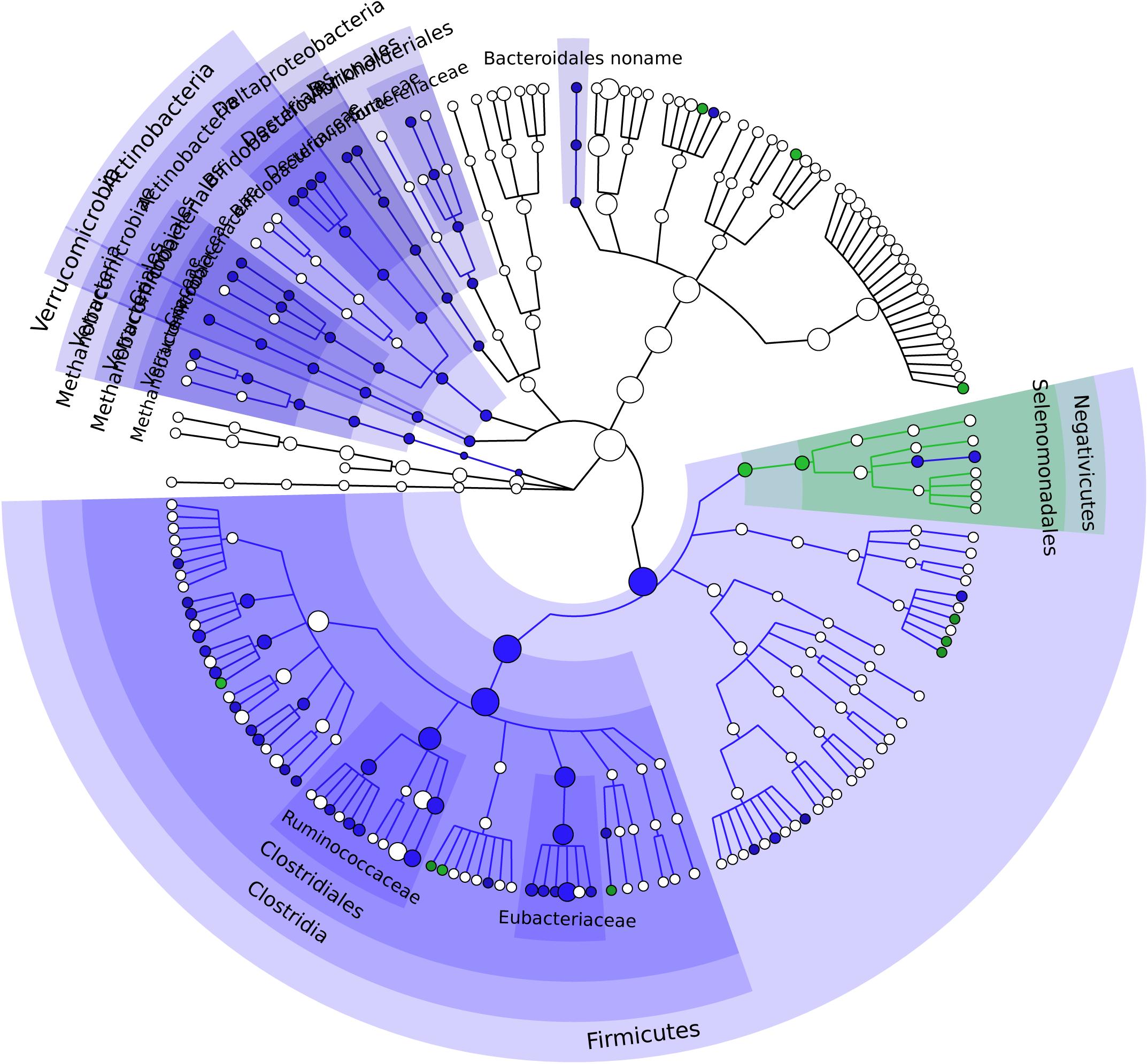
Differences in intestinal microbiota taxonomic structure before and after *Helicobacter pylori* eradication therapy (based on LEfSe analysis with default parameters). GraPhlAn cladogram (circular hierarchical tree) of samples grouped by 1st and 2nd time points. Each dot represents bacterial taxon. OTUs significantly increase in 1st time point shown by blue color. Similarly, OTUs significantly increase in 2nd time point denoted by green color. The various shades of colors highlighting the background indicate that the changes consistent at several taxonomic levels.

We compared the alpha-diversity and the Shannon indices between the 1st and the 2nd time points. The alpha-diversity of the intestinal microbiota composition differed significantly between the metagenomes before and after the eradication therapy: for the observed species Mann-Whitney P=2.80×10^−5^, for Jensen-Shannon index P=0.01 (see Supplementary Figure S3). However, there was no clear evidence for difference between the metagenomes of the 1st and 2nd time points based on the taxonomic composition (ANOSIM of Bray-Curtis dissimilarity values, R=0.075, P=0.002, 10 000 permutations). Distribution of the patients’ gut community structures compared to the healthy persons (Voigt et al., 2015) at two time points is shown in Figure 2.

**Figure 2.**
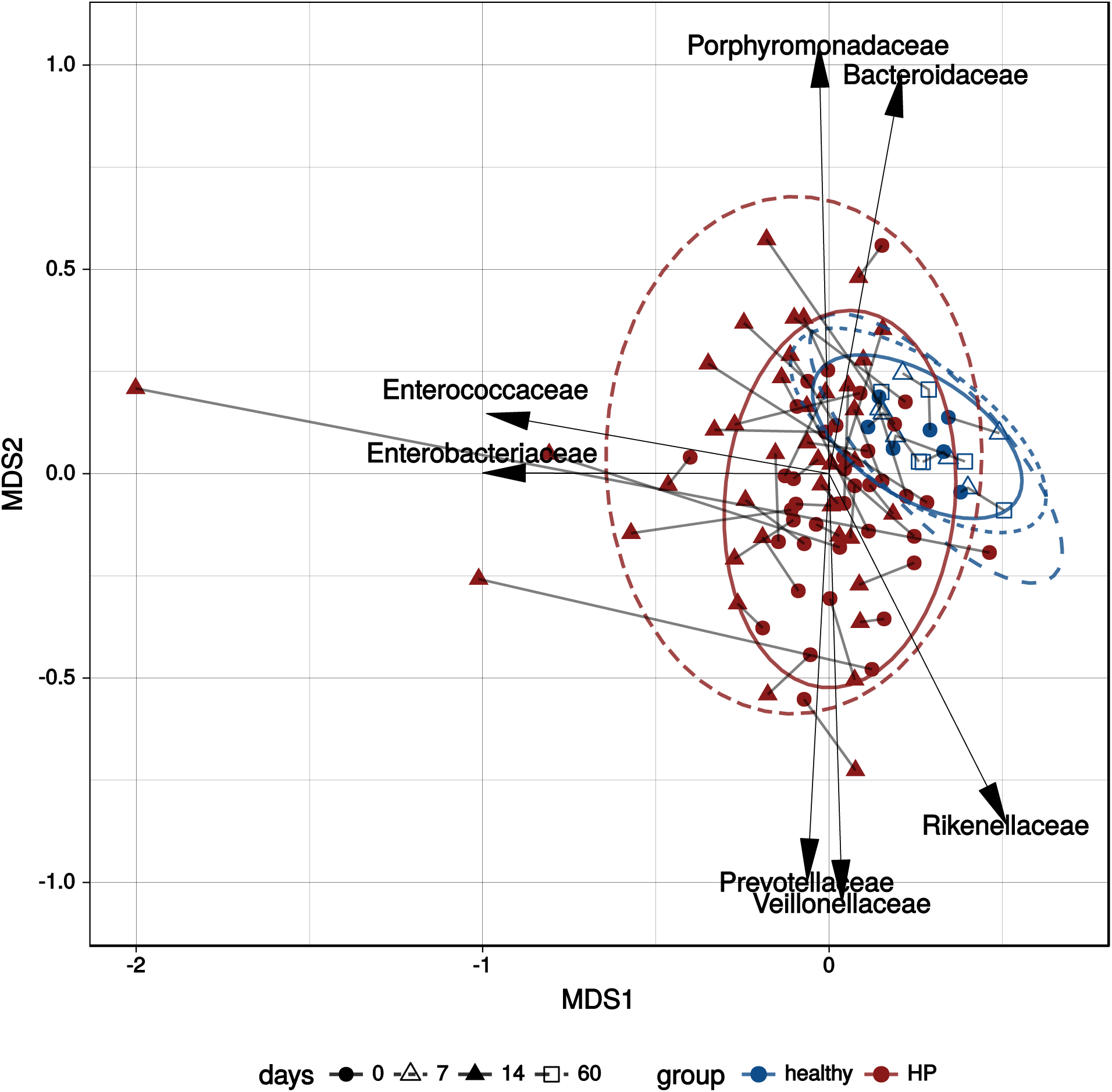
MDS of gut microbial community structure of the patients prior to and after therapy compared to the microbial community structure changes of the healthy individuals at different time points (Voigt et al., 2015). Multidimensional scaling (MDS) biplot using Bray-Curtis dissimilarity metric. The labels denote the directions of the increasing abundance of the respective microbial family. Red color denotes patients with *H. pylori* eradication group, blue color refers to the control one. Time points are shown by marks of different shape. The samples collected from the same patient are connected by lines. The components have parameters P<0.05, R^2^ >0.15 according to envfit analysis from the R vegan package (Oksanen et al., 2015).

To visualize the scale of changes in the taxonomic profile, the change in the alpha-diversity (Shannon / Simpson index) and the Bray-Curtis distance between the metagenomes of each patient at the 1st and 2nd time points was plotted (Figure 3). We observed that some metagenomes of the experimental group have undergone moderate (below mean value), medium (above mean value) and severe (higher than mean+SD) changes, whereas the metagenomes of those individuals who didn’t take therapy (Voigt et al., 2015) did not show severe changes in the taxonomic composition on days 7 and 60 and were in the moderate zone. There were statistically significant differences in Bray-Curtis distance between time points in the experimental and control samples: Mann-Whitney P = 2.43 × 10^−9^.

**Figure 3.**
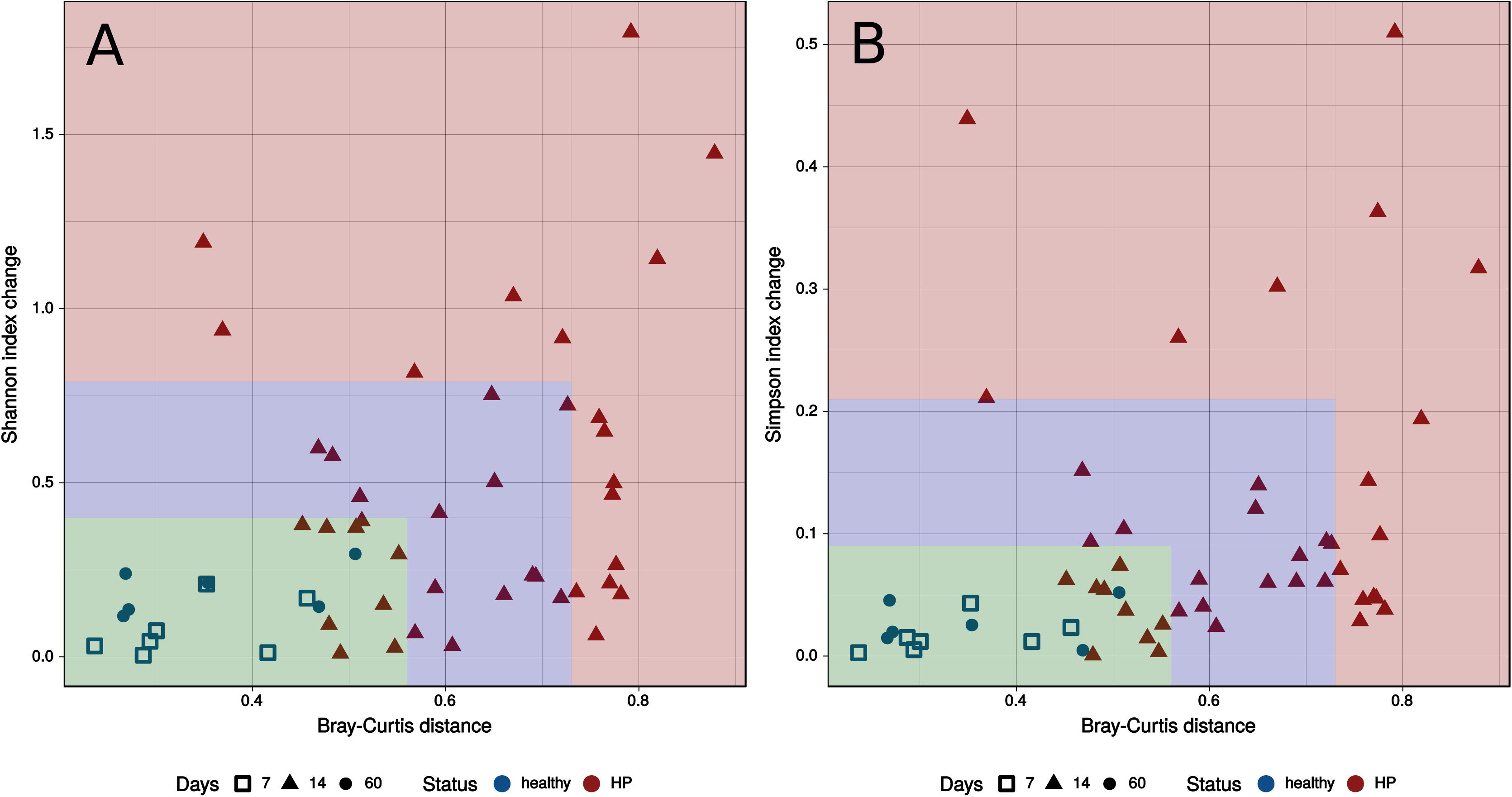
The scales of changes in the gut microbiota taxonomic composition caused by *H. pylori* eradication therapy. The X-axis shows the Bray-Curtis distance (**A, B**) between the 1st and 2nd time points. Y-axis denoted the module of difference between the alpha diversity indexes such as Jensen-Shannon (**A**) and Simpson (**B**) of the 1st and 2nd time points. Case group variables are shown as the red dots, the blue dots denote control group variables (Voigt et al., 2015). The green color area show change below mean value (<mean), blue color area - above mean value (>mean, but <mean+SD), variables above mean value are in the red color area (>mean+SD).

Conclusions, based on the taxonomic profiles obtained using MetaPhlAn2, were confirmed by MetaFast which is taxonomic-free computational method based on the *de novo* combined adaptive assembly of metagenomic reads (Ulyantsev et al., 2016). The results are shown in the Supplementary Figure S4. Dependence of the metagenomes distance of the 1st time point from the control group and the scale of microbiota changes after eradication therapy were not identified (Spearman R=0.20, P=0.21, see Supplementary Figure S5A). However, the greater the change after the therapy, the farther the 2nd time point from the control group (Spearman R=0.64, P=1.4×10^−5^, see Supplementary Figure S5B). On the other hand, the scale of changes associated with *H. pylori* eradication therapy may depend on some specific characteristics. The study of the different clinical factors effect on the patient’s intestinal microbiota taxonomic composition is given below. The relative abundances of microbial genera/species in control groups are shown in Supplementary Tables S4 and S5.

### The power of *H. pylori* eradication therapy effect on gut microbiota is different depending on disease and gender

Analysis of variance using distance matrix (ADONIS of Bray-Curtis dissimilarity, 10 000 permutations) was performed to reveal factors which influence microbiota taxonomic composition. According to the obtained results, the most relevant factors shaping microbiota composition are gender (R^2^=3.97, adj. P=0.0027), time point (R^2^=3.53, adj. P=0.0036), and duodenal ulcer (R^2^=2.03, adj. P=0.0836). It should be noted that patients with duodenal ulcer have shown more significant distance change from the 1st time point compared with the rest of the patients (Mann-Whitney P=0.0076). Similar effect was observed in female patients regardless of diagnosis (Mann-Whitney P=0.0050) (see Supplementary Figure S6).

The significant factors revealed were further used in MaAsLin analysis workflow. It was shown that relative abundance decrease of 7 genera and 13 species was associated with eradication therapy treatment (q-value<0.05). The taxa with decreased abundance include Bifidobacterium (*Bifidobacterium adolescentis, Bifidobacterium longum*), Eubacterium (*Eubacterium biforme, Eubacterium ventriosum*), Ruminococcus (*Ruminococcus callidus, Ruminococcus lactaris*), *Roseburia hominis*, *Collinsella aerofaciens*, *Dorea longicatena, Akkermansia muciniphila, Bacteroidales bacterium ph8, Dialister invisus, Coprococcus catus*. The data is shown in Supplementary Figure S7. For factors gender and duodenal ulcer, changes were determined with 0.25>q-value>0.05.

The results of the MaAsLin analysis present in GitHub repository (https://github.com/OlekhnovichEvgenii/HP-project/).

### Functions of gut microbiota after *H. pylori* eradication therapy propensity toward metabolic profile perturbation and increase macrolide resistance potential

We estimated the functional capacity of patients’ gut microbiota in different time points. Following eradication treatment seven KEGG pathways associated with cellular processes (ko02030 - bacterial chemotaxis, ko02024 - quorum sensing, ko02040 - flagellar assembly), genetic Information processing (ko00970 - aminoacyl-tRNA biosynthesis and ko03010 - ribosome) and environmental Information processing (ko02010 - ABC transporters and ko02020 - two-component system) were increased after eradication therapy. Simultaneously, five KEGG metabolic pathways associated with metabolism of cofactors and vitamins (ko00130 - ubiquinone and other terpenoid-quinone biosynthesis and ko00785 - lipoic acid metabolism) and glycan biosynthesis and metabolism (ko00511 - other glycan degradation and ko00540 - lipopolysaccharide biosynthesis) were decreased. It is worth noting the decline in overall metabolic potential (ko01100 - metabolic pathways decreased). Results are present in Supplementary materials (see Supplementary Tables S6-S7).

To study human gut microbiota functional shift in more detail, we analysed three specific microbiota functions which may be associated with eradication therapy components. Firstly, we studied antibiotic resistance as antibiotic treatment may have influence the distribution of ARGs in the microbial community (Modi et al., 2014). The second microbial community function under study was resistance to biocide and heavy metals because eradication therapy includes bismuth subsalicylate treatment. As a third characteristic, we considered virulence factors as it has been shown that following antibiotic treatment opportunistic microorganisms get an advantage (Modi et al., 2014).

We have revealed 135 genes groups which determine resistance to 16 antibiotic classes (Supplementary Tables S8). ARGs amount were increased following eradication therapy (Mann-Whitney P=0.006). Relative abundance of ermB (23S rRNA methyltransferase) gene associated with macrolide resistance increased significantly (adj. P=0.003). The number of SNPs in the ermB gene have also increased after the therapy (Mann-Whitney P=0.03), which may be a circumstantial evidence of the increase in the number of the microorganisms that possess this gene. At the same time, the functions which determine tetracycline resistance were decreased (adj. P=0.019) (see Supplementary Table S9). Changes in the abundance of the resistance genes to biocide/metals and virulence factors were not detected (mapping results are present in Supplementary Tables S10-S11).

### *H. pylori* eradication therapy affects the distribution of intraspecies structures

To analyze microbiota composition in more detail, i.e. on the intraspecies structure level, we used ConStrains. To detect intraspecies structures, this approach uses SNPs patterns found in the set of universal genes (Luo et al., 2015). Totally, 500 strains (which is the sum of unique strains for each patient) were detected in the experimental group at the 1st and 2nd time points. We identified 371 intraspecies structures (total sum of the unique strains for each healthy individual) that formed a control sample. Intraspecies structure composition before and after treatment have been compared. The data obtained is shown in Figure 4. It was shown that gut microbiota intraspecies structures within the case group are biased to scatter more than those of the control group. Bray-Curtis distance scales between the two groups was found to be significant (Mann-Whitney P=0.01).

**Figure 4.**
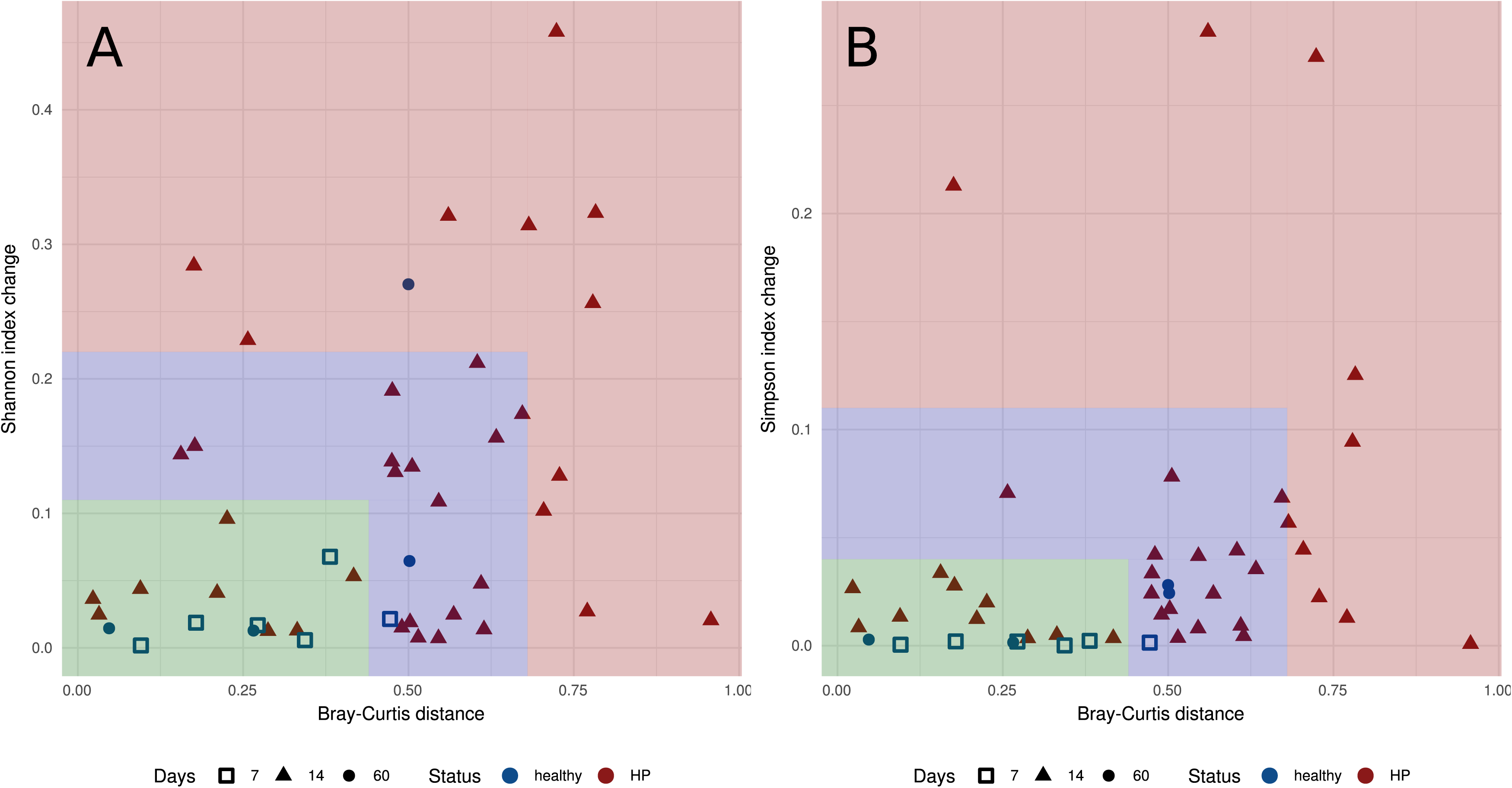
Intraspecies changes of observed “strains” relative abundance in case and control groups (Voigt et al., 2015). The X-axis shows the Bray-Curtis distance (**A, B**) between the 1st and 2nd time points. The Y-axis denoted the module of difference between the alpha diversity indexes such as Jensen-Shannon (**A**) and Simpson (**B**) of the 1st and 2nd time points. The variables of case group are shown as the red dots, the blue dots denote control groups. The green color area show change below mean value (<mean), blue color area - above mean value (>mean, but <mean+SD), variables above mean value denoted red color area (>mean+SD).

We have compared the relative abundance of each gut microbiota intraspecies structure within the case and control groups. Only structures with nonzero relative abundance in two time points were included into the analysis. In control group metagenomes sampled following 7 and 60 days after the 1st sample collection were analyzed separately. The data is presented in Supplementary Figure S8. The analysis revealed that intraspecies structures relative abundance in the experimental group is more unstable than that of the control group. We observed several species which relative abundance was almost constant but strains distribution varied greatly (i.e. *Faecalibacterium prausnitzii*, *Prevotella copri*, *Alistipes putredinis*, *Bacteroides vulgatus, Bacteroides uniformis* - see Supplementary Figure S9).

The PanPhlAn was used to study the functional capacity of specific bacterial species using microbial community data. This strain-level metagenomic profiling tool aims to identify pangenomes of individual strains in metagenomic samples (Scholz et al., 2016). We were able to extract specific genes content from metagenomes for the most abundant taxa in the experimental group. To perform the analysis, two taxa (*E. rectale* и *F. prausnitzii*) were selected and their gene compositions were compared at different time points. The changes on KEGG Orthology (KO) groups abundances for both *F. prausnitzii* and *E. rectale* strains were found (see Supplementary Tables S12 and S14). For *E. rectale*, 212 KOs increased after antibiotics therapy, 17 - decreased; for *F. prausnitzii* 5 KO increased and 30 KO - decreased. We merged KO by KEGG pathways and counted the number of changing genes for each pathway (see Supplementary Tables S13 and S15).

It should be noted, the observed changes in the metabolic profile of *E. rectale* were more extensive (this may be due to a higher reads coverage in the metagenomes of this bacterium, which allowed to extract more representative gene content distributions). The genes associated with ko01110 (biosynthesis of secondary metabolites), ko01130 (biosynthesis of antibiotics), ko02040 (flagellar assembly), ko02010 (ABC transporters), ko02030 (bacterial chemotaxis), ko02020 (two-component system) were increased that could be useful for survival in changed environmental conditions.

These species in control group don’t showed significant changes (threshold adj. P < 0.01).

### *H. pylori* eradication therapy affects strains distribution of *Enterococcus faecium* within microbiota

*Enterococcus* spp. multiple antibiotics resistance is alarming for the world medical community as this bacterium is prevalent among clinically significant microorganisms. Enterococci may collect different mobile elements and spread antibiotic resistance factors to the broad spectrum of gram-positive and gram-negative bacteria, including the transfer of vancomycin resistance to methicillin-resistant strains of *Staphylococcus aureus* (Lebreton et al., 2017).

We managed to recover *E. faecium* at two time points from the stool samples of only two patients (HP003, HP010). Strains isolated after the therapy had increased resistance to antibiotics (see Supplementary Table S16), and had higher numbers of ARGs to macrolide and tetracycline antibiotics (ermB, tetL, tetM genes) (Supplementary Figure S10A). Interestingly, *E. faecium* strains isolated after the therapy (*E. faecium* of 2nd time point) from both patients were similar in the SNPs pattern, and differed from *E. faecium* strains isolated at the 1st time point (Supplementary Figure S10B). Also, *E. faecium* of 2nd time point relative abundance in the metagenomes after the therapy has grown when compared to the 1st time point isolates (Supplementary Figure S10C and Supplementary Figure S10D). A more detailed description of the results is given in the supplementary materials (Supplementary Note 1). Also, both strains were close with gut commensal [Bender et al., 2015], surrogate (using in place of pathogens for validation of thermal processing technologies and systems) [Kopit et al., 2014] and clinical *Enterococcus faecium* strains [Mbelle et al., 2017; Sassi et al., 2018] (BLAST results present in Supplementary Table S17).

### Metagenomic context of the ermB gene shows potential localization in genomes of distant bacterial taxa

To study in more details the process of acquiring ARGs, the metagenome samples of patients HP003, and HP010 were re-sequenced with Illumina HiSeq 2500. It allows to obtain paired reads of the length sufficient for metagenome assembly. Using MetaCherchant tool (see Materials and methods) we performed the target assembly of the ARGs genomic context. Based on the topology of the obtained graph, we established that the ermB gene was present in *Enterococcus* spp. and *Bacteroides* spp. genomes (Figure 5). The relative abundance of these bacteria increased after the eradication therapy (see Supplementary Figure S11). Also worth noting is the co-localization of the ermB gene with the tetL and tetM genes coffering the resistance to tetracyclines.

**Figure 5.**
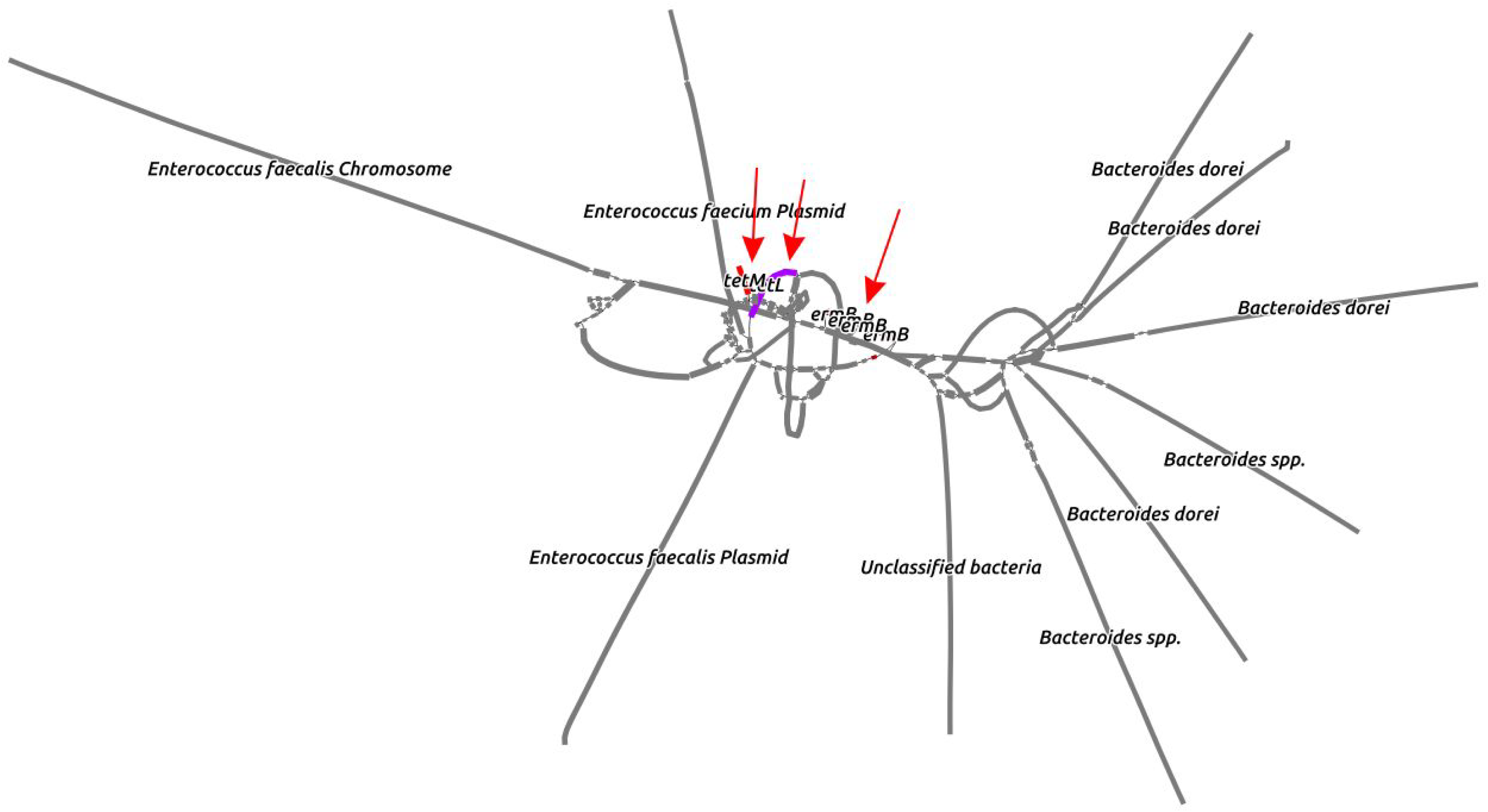
The graph of ermB gene context obtained by MetaCherchant from HP003 patient (2nd time point) metagenomic data. The resulting graph showed target gene co-localization with other ARGs - tetL and tetM. ARGs denoted by red arrows. Taxonomic annotation is shown according to the BLAST results.

One of the possible scenarios may be that certain nosocomial strain of *E. faecium* colonized the HP003 patient’s intestine (because it is identical in the pattern of SNPs in another patient). It carried the mobile element containing the ARG of resistance to macrolides (ermB), which allowed to gain an advantage and grow in the relative abundance during the antibacterial therapy to the donor itself (*E. faecium*) and recipients - *Bacteroides dorei* and *Enterococcus faecalis*. On the other hand, the ermB gene could be present in the patient’s metagenome before the therapy, but the reads coverage was insufficient to detect its presence (the context of the ermB gene at the 1st time point for patient HP003 was not assembled due to insufficient coverage) and *E. faecium* acted as a recipient of this function. The third possible scenario for the development of events may be the assumption that the genes of ermB *Enterococcus* spp. and *Bacteroides* spp. are independent homologues localized in mobile genetic elements containing homologous sequences.

For the HP010 patient, it was not possible to reconstruct the environment of the ermB gene for any time point due to low coverage.

## Discussion

Changes in the microbial community structure caused by *H. pylori* eradication therapy have been the subject of study before. Yap et al. (2016) used 16S sequencing technique to study microbiota in young adults after eradication treatment. The authors showed that 6 months following the treatment Firmicutes relative abundance decreased, and Bacteroides abundance increased. However, the authors summarize that in general microbial community remains unchanged which may be due to the rather long time period that passed since the treatment compleet and compensatory processes that took place in the microbiota community.

Gut microbial community structure is altered by antibiotic treatment, which was shown both by the results described in this article and the data obtained by other investigators (Willmann et al., 2015; Korpela et al., 2016). We showed that eradication therapy decreases the wealth of microbial community. It should be noted that patients with duodenal ulcer and female patients have shown more significant changes in the microbial community taxonomy structure. For female patients, this observation could be explained by a different phase of the menstrual cycle in which samples were taken. On the other hand, microbiota of female individuals from the control group didn’t show wide shifts within 60 days. It appears that patients with duodenal ulcer showed significant changes due to their reduced condition. Possibly, duodenal ulcer is most important condition for gut microbiota in comparison with stomach ulcer / gastritis.

We have revealed taxonomy change biomarkers which appear to be associated with eradication therapy. We observed the decrease of *B. adolescentis* and *B. longum* that are involved in the carbohydrate metabolism and prevent colonization of the gut mucosa by opportunistic species (Hütt et al., 2006; Martins et al., 2010), of *C. aerofaciens* associated with decreased risk of inflammatory bowel disease (Kassinen et al., 2007), and of *A. muciniphila* which abundance reduction is thought to be associated with increased weight (Everard et al., 2013; Dao et al., 2015; Plovier et al., 2017). Abundance decrease in some other species, such as *D. longicatena, Bacteroidales bacterium ph8, D. invisus, C. catus, E. biforme, E. ventriosum*, *R. callidus, R. lactaris*, *R. hominis* have also been found. Proton pump inhibitors may cause intrusion of the gut microbiota community by the bacteria species which can’t pass gastric acid barrier under normal condition (Jackson et al., 2016; Imhann et al., 2016). However, we didn’t reveal any tendency which agrees with this proposition.

We determined statistically significant differences in microbiota functional profiles between the 1st and the 2nd time points. In general, the decline in seven KEGG pathways associated with cellular processes (ko02030, ko02024, ko02040), genetic Information processing (ko00970, ko03010) and environmental Information processing (ko02010, ko02020) were increased after eradication therapy. Simultaneously, five KEGG metabolic pathways associated with metabolism of cofactors and vitamins (ko00130, ko00785) and glycan biosynthesis and metabolism (ko00511, ko00540) were decreased. The increasing pathways may be associated with better survival potential in the changed environment. Interestingly, we determined decline in ko01100 pathway (Metabolic pathways), this may be due to a overall impoverishment of the gut community.

We also detected prevalence of tetracycline ARGs in patients’ microbiota before the therapy, which is consistent with previous findings (Hu et al., 2013). However, these genes decreased after therapy. Genes that determine resistance to macrolides (ermB, 23S rRNA methyltransferases group) were found to increase at the second time point. Similar results showing an increase of genes belonging to the 23S rRNA methyltransferases group were reported by Jakobsson et al. (2010). It should be noted, the specific and non-specific mechanism of resistance to antibiotics were increased after eradication therapy.

Microbiota serves as ARGs reservoir and can transmit these genes to pathogenic bacteria. The decrease in relative abundance of the tetracycline resistance factors is of interest. This effect could be caused by the decline of the bacteria which possess these genes, as tetracycline is not included in the eradication therapy, and presence of the genes which determine resistance to this antibiotic doesn’t give the critical advantage to their carriers under given circumstances. Genes which determine resistance to biocide, heavy metals, and also virulence factors remained unchanged.

Interestingly enough, changes of the intraspecies structure distribution were more prominent in the experimental group compared to the control one. We observed perturbations in strains abundances even for those species which abundance were almost constant. This result stresses an importance of strain profiling in metagenomic research as different strains of bacteria can vary significantly in phenotype. The changes in the metabolic profile of *E. rectale* were observed. The genes associated with ko01110 (biosynthesis of secondary metabolites), ko01130 (biosynthesis of antibiotics), ko02040 (flagellar assembly), ko02010 (ABC transporters), ko02030 (bacterial chemotaxis), ko02020 (two-component system) were increased that could be useful for survival in changed environmental conditions. These results may be compared with changes on overall metabolic profile.

It seems that strains carrying ARGs, i.e. 23S rRNA methyltransferases, gained an advantage under the strong selective pressure. The *E. faecium* strains isolated from two patients after eradication therapy treatment have shown an increased antibiotic resistance *in vitro*. These strains had enhanced resistance to macrolides, which is determined by 23S rRNA methyltransferase gene, and possessed virulence factors, such as collagen adhesin protein Scm. In addition, these strains were more abundant after the treatment when compared to the strains isolated at the first time point. These observations are consistent with the findings of the bioinformatics analysis.

Horizontal gene transfer (HGT) could be another possible way which enhances microbiota resistance potential to antibiotics. Using MetaCherсhant tool, we constructed the graph structure which contains members of *Enterococcus* and *Bacteroides* spp. and represents metagenomic context of the ermB gene. This graph structure clearly shows that the target ermB gene is localized in close proximity to the tetracycline ARGs (tetL and tetM). It can be assumed that these resistance factors can spread together and cause multiple antibiotics resistance by HGT. The result obtained could be a circumstantial evidence of the ermB gene spread within the intestinal microbiota. This finding certainly requires additional verification, but it shows that new computational methods are able to predict such events.

Thus, *H. pylori* eradication therapy causes multiple shifts and alterations of the intestinal microbiota structure and leads to the accumulation of genes which determine resistance to macrolides. It is worth noting, microbial community demonstrated shift to reduce metabolic potential and to increased mechanisms, which mediate more survival condition through intraspecies perturbations. Study of antibiotic treatment in terms of its impact upon the human gut microbiota allows shedding light on of the complex processes that cause accumulation and spread of antibiotic resistance. An identification and understanding of these complicated processes may help to constrain antibiotic resistance spread, which is of great importance for human health care.

## Materials and methods

### Sample collection

Faecal samples were acquired prior to and following eradication therapy from the patients suffering from diseases associated with *H. pylori* infection - gastroesophageal reflux disease (GERD), chronic gastritis, chronic duodenitis, and duodenal ulcer (see Supplementary Table S1). In all cases the *H. pylori* infection was confirmed by urease testing and cytological examination of mucosal biopsies collected from the antral part of the stomach during gastroscopy study. All patients included in the study received the *H. pylori* eradication treatment according to the Maastricht scheme (Malfertheiner et al., 2017), which included amoxicillin 1000 mg, clarithromycin 500 mg, bismuth subsalicylate 240 mg, proton pump inhibitor (esomeprazole / pantoprazole) 20 mg. The therapy course lasted for 14 days. Lactulose was used as a prebiotic throughout the therapy course. Before the start of the study, each patient signed an informed consent. Two sampling points were included: before the treatment and 0 or 2 days after the treatment. A total of 80 fecal samples from 40 patients were collected (47.7 ± 13.11 years old, 18 female and 22 male).

### Sample preparation and sequencing

Stool samples were delivered to the laboratory and subsequently were stored at - 20 °C. Before the DNA extraction procedure, the aggregate state of the samples was evaluated and their weight was determined. The weight of all the samples exceeded 20 grams, which was a sufficient quantity for the extraction of the required amount of DNA. Total DNA extraction was performed as described previously (Tyakht et al., 2013). The whole genome shotgun sequencing on SOLiD 5500W platform in the 50 base pairs in single-end mode was performed according to the manufacturer’s recommendations (Life Technologies, Foster City, CA, USA) using the following kits: 5500W Conversion Primers Kit, 5500 W FlowChip V2, 5500 W FlowChip Prep Pack, 5500W Template Amplification Kit v2, 5500 W FWD1 SP Kit, Double, 5500W FWD2 SP Kit, Double, 5500W FWD SR Kit, Double, 5500W FWD Ligase Kit, Double, 5500W Run Cycle Buffer Kit, 5500W FWD Buffer, Double, 5500W Buffer D.

### Additional experimental data

The dataset from longitudinal study of gut metagenomes without intervention was used for the comparative data analysis and visualization (Voigt et al., 2015). Also, we used previously published data from metagenome projects - ENA project ID PRJEB21338 (Boulygina et al., 2017), ENA project ID PRJEB18265 (Gluschenko et al., 2017), and from the NCBI bioproject ID PRJNA412824 (Prianichnikov et al., 2018) harboring genomes of *Enterococcus faecium* isolates. The methods of *E. faecium* isolation and susceptibility testing are described in Supplementary Note 2. Summary information on the samples used in the study is available in the Supplementary materials (Supplementary Tables S18-S20).

### Metagenomic sequences data preprocessing

The metagenomic reads were filtered by quality. Reads with an average quality value (QV) lower than 15 were discarded. To minimize sequencing errors, the remaining reads were processed with SAET software (Life Technologies) with parameters -qvupdate -trustprefix 25 -localrounds 3 -globalrounds 2. Further quality filtration was carried out as following: all positions starting from 5’-end were removed up to the 1st high quality position (QV ≥ 30). After filtering all the reads with the length less than 30 nucleotides were discarded. In order to remove fragments of human DNA from the data, the reads were mapped to the human genome (Homo sapiens hg38) using Bowtie (Langmead et al., 2009) with parameters -f -S -t -v 3 -k 1. If a read could be mapped in two or more places, the reference was chosen randomly and equiprobable. Reads unmapped to the human genome were converted from the color space to base space and used in further analysis.

### Taxonomic and functional analysis of the metagenomes

Metagenomic taxonomic and metabolic profiling was performed using HUMAnN2 (Abubucker et al., 2012) (contains MetaPhlAn2 (Truong et al., 2015)) using Kyoto Encyclopedia of Genes and Genomes (KEGG) database (Kanehisa et al., 2015). ARGs were identified in the metagenomes by mapping the reads to MEGARes database (Lakin et al., 2016) using Bowtie with keys -f -S -t -v 3 -k 1. Relative abundance of the group and class ARGs was searched using ResistomeAnalyzer (Lakin et al., 2016) and calculating RPKM (reads per kilobase per million mapped reads) value:

> *num reads / ( gene length/1000 * total num reads/1 000 000),*

where *num reads* - the number of reads mapped to a gene sequence, *gene length* - length of the gene sequence, *total num reads* - total number of mapped reads of a sample. The number of SNPs in ARGs was revealed by SNPFinder (Lakin et al., 2016). MetaCherchant was applied for the specific metagenomic assembly of the target ARGs context (Olekhnovich et al., 2018). The taxonomic annotation of resulting graph nodes was performed using BLAST.

For each ARGs, resistance gene (R-genes) to biocides and heavy metals, as well as the virulence factors, homologous genes in the 9.9 mln catalog of human gut microbiota genes (Qin et al., 2010) were determined using BLAST (homology criterion: e-value < 10^-5^, percent identity > 80% for > 80% of length) and two databases: BacMet (Pal et al., 2013) and VFDB (Chen et al., 2012). The relative abundance of each template gene was calculated by summing the values of the number of such similar genes from the above-mentioned catalog (according to Dubinkina et al. (2017)). RPKM value was calculated to normalize the mapping data.

Distribution of intraspecies structures in metagenomic data was obtained using ConStrains (Luo et al., 2015) with default parameters. The scale of changes in the relative abundance of intraspecific structures was calculated as a residual of abundances at two time points. The pangenomes of the most abundant bacteria were obtained by PanPhlAn (Scholz et al., 2016) with parameters --left_max 1.5 --right_min 0.5. The functional annotation of pangenomes (gene content) was performed according to KEGG Orthology (KO) (Kanehisa et al., 2015).

### Statistical analysis

The basic identification of differences at a taxonomic level was performed using linear discriminant analysis effect size (LEfSe) (Segata et al., 2011). The differences between community structures were evaluated using the Bray-Curtis metric. For additional analysis of the metagenomic composition, the MetaFast algorithm based on the adaptive assembly of *de novo* metagenomic reads was used. This algorithm evaluates the differences in metagenomes without the use of taxonomic annotation methods (Ulyantsev et al., 2016). The ADONIS with the Bray-Curtis dissimilarity metric was used to identify the factors affecting the patients microbiota taxonomic composition (Oksanen et al., 2015). Factors included gender, age, duodenal ulcer, chronic duodenitis, chronic gastritis, GERD and time point. The factors which significantly (threshold P<0.1) influenced the microbiota taxonomic composition were added to the multivariate analysis MaAsLin (Morgan et. al., 2012). Relative abundance of different species and genera were considered as dependent variables. Analysis scheme was as follows:

> *relative abundance of genus/species∼time point+duodenal ulcer+gender*

To determine the differences in relative abundance of KEGG metabolic pathways, ARGs, resistance genes to biocide / metals and virulence factors the following operation were performed: the differences in the representation profiles were determined using the paired Mann-Whitney test (significance threshold: adjusted P<0.01) then significant differences of features were identified using gene set analysis (GSA) from piano bioconductor package (Väremo et al., 2013) (parameters: gene set analysis using “reporter feature algorithm”, significance assessment using “gene sampling”, gene set significance threshold: adjusted P<0.05 (Dubinkina et al. 2017). Differences in KO abundances of extraction pangenomes by PanPhlAn was performed using Fisher exact test with Benjamini-Hochberg correction for multiple testing (FDR) (significant threshold adj. P<0.01).

Paired Mann-Whitney test was used for the additional differential analysis. Data visualization was performed using R (R core team, 2016) and GraPhlAn (Asnicar et al., 2015).

## Availability of data and addition materials

Raw metagenomic reads for fecal samples are deposited in the NCBI Archive (project ID: PRJNA413659; Web-address: https://www.ncbi.nlm.nih.gov/bioproject/413659). Scripts used in this work and other additional materials are available at GitHub (https://github.com/OlekhnovichEvgenii/HP-project).

## Ethics approval and consent to participate

The study was approved by the ethical committee of the Kazan State University. Before the start of the study, each patient signed an informed consent.

## Acknowledgements

This study were financially supported by the Russian Scientific Foundation (grant # 15-14-00066). Prof Kruglov AN for bacteriological testing of stool samples.

## Competing interests

The authors declare that they have no competing interests.

